# Phylogenetic evidence for the emergence of bacterial σ-factors by domain fusion

**DOI:** 10.64898/2026.10.08.757525

**Authors:** Pragya Tiwary, Aswin Sai Narain Seshasayee

## Abstract

σ-factors are dissociable subunits of RNA polymerase (RNAP), forming the basal regulatory machinery for bacterial transcription. The minimal σ-factors contain two domains (called σ_2_ or σ_4_), both of which bind RNAP as well as the promoter DNA through their helix-turn-helix motif. Housekeeping σ-factors contain four domains. We identify proteins containing at least one of the four σ-factor domains in bacterial and bacteriophage genomes; we also show that proteins containing either σ_2_ or σ_4_, which are not defined as σ-factors, are commonly found in both sets of organisms. Using phylogenetic analysis of these two domains, we suggest that the highly conserved major housekeeping σ-factor, with its complex domain architecture, represents a late-evolving monophyletic clade, whereas single-domain proteins and minimal σ-factor architectures comprising σ_2_ and σ_4_ might be ancestral. Thus, σ-factors probably evolved from single-domain DNA-binding transcription factors by domain accretion.

## Introduction

All multisubunit RNAPs comprise a core enzyme and dissociable specificity subunit(s). In bacteria, these specificity subunits are called sigma factors (σ-factors) [1], [2]. σ-factors are essential for recognising specific DNA promoter sequences, positioning the RNAP holoenzyme at the target promoter, and enabling the local unwinding of the DNA duplex near the transcription start site (TSS) [3]. The σ70 family usually initiates transcription at ™35/™10 promoters [4], and the σ54 family requires specific activators to start transcription at ™24/™12 promoters [5]; these are the two evolutionarily and structurally distinct families of σ-factors. Along with numerous other σ70 and/or σ54 factors that regulate more specialised gene sets (such as sporulation and heat shock), all bacteria encode a principal house-keeping σ70 factor that controls the majority of transcription during exponential growth. The number of σ-factors encoded varies across bacteria; for instance, *E. coli* encodes seven σ-factors, *Bacillus subtilis* has eighteen [6], while *Streptomyces coelicolor* has more than sixty σ-factors [7]. σ-factors have also been detected in bacteriophage genomes [8]; a few have been experimentally characterised [9] and these appear to help the bacteriophage manipulate the host transcriptional machinery for its own benefit [10].

The σ70 family has been divided broadly into four groups based on domain architecture. Group 1 consists of the housekeeping σ-factors (RpoD in *E. coli*), having four domains, conventionally numbered 1-4 (and referred to here and elsewhere as σ_1_, σ_2_, σ_3_, and σ_4_). Group 2 σ-factors are dispensable proteins with domain architecture similar to that of group 1 (but lacking the N-terminal 1.1 region of σ_1_). Group 3 σ-factors contain domains σ_2_, σ_3_, and σ_4_ (RpoH in *E. coli*) [11]. Finally, group 4 accommodates the numerically largest extra-cytoplasmic function (ECF) subfamily; these are the ‘minimal size’ σ-factors having only domains σ_2_ and σ_4_, connected via a highly disordered and poorly conserved linker (RpoE in *E. coli*) [12]. We define σ-factors as basal regulators of transcription to distinguish them from transcription factors (TFs).

TFs are proteins that usually bind to the DNA upstream or around the promoter and repress or activate transcription [13], [14]. Transcription activation typically involves the TF presenting an interface to recruit the RNAP after the TF has bound to the DNA [15]. In contrast, σ-factors bind to the core, catalytically competent RNAP, to form a ‘holoenzyme’, which then specifically binds promoters [16]; the binding of the holoenzyme to promoters can be modulated by TFs. Bacteria use a combination of σ-factors and TFs to regulate gene expression.

The evolution of TF sequences and their repertoire under a phylogenetic framework has been explored in recent times in both prokaryotes and eukaryotes. Visweswariah and Busby have argued that functions that typify TFs, such as sequence-specific DNA binding, oligomerisation, RNAP interfaces (in the case of activators), and signal sensing, might have evolved atop an ancestral non-sequence-specific DNA-shaping protein [17]. An example of such evolution is the HU/IHF family of DNA-binding proteins, in which the sequence-specific IHF might have evolved from a non-specific HU-like ancestor [18]. We, however, lack information on how basal regulators such as σ-factors evolved in the first place. We address this question for σ-factors belonging to the σ70 family, which are encoded in both bacterial and bacteriophage genomes. Using a dataset of over 60,000 σ-factor-like protein sequences, we analyse how the different domains that characterise σ-factor classes have evolved and how they might have fused or dissociated to form σ-factors as we know them.

## Materials and Methods

### Dataset Curation

A list of 873,718 phage proteomes was downloaded from the PhageScope database [19]. Redundancy was removed by filtering through average nucleotide identity (ANI) [20], resulting in a final dataset of 351,065 proteomes. The bacterial dataset comprised 1,444 genus-level non-redundant proteomes, used in a previous study [18].

### Identification of sigma-70 proteins

To find domain homologs, the hmmsearch command from the HMMER suite [21], [22], [23] was run with a trusted cutoff threshold for the PFAM profiles [24] of sigma factor domains (PF08281, PF04542, PF04539, PF04545, PF00140, PF03979, PF04546, PF07638, PF04552, and PF04963) against the bacterial and phage databases separately. The search produced a total of 32,488 phage and 28,909 bacterial sigma proteins.

### Phylogenetic Tree Construction

A final set of 22,084 (3,003 phage and 19,081 bacterial) σ_2_ sequences and 45,221 (7,412 phage and 37,809 bacterial) σ_4_ sequences was used to make an alignment. The alignment was done using a) Muscle v5 [25] and b) hmmalign from the HMMER suite [23]. For σ_4_ phylogenies, hmmalign was done separately using the hits pertaining to two different PFAMs (PF08281 and PF04545). For Muscle-aligned sequences, the resulting alignment was trimmed using trimAl with a 0.02 cutoff to remove excessive gap-containing columns [26]; however, no trimming was done for the HMM-aligned sequences. These alignments were then subsequently used to build approximate maximum-likelihood phylogenetic trees using FastTree 2.2 [27], wherein the model used was the WAG (Whelan-And-Goldman 2001) model [28].

The full set of sequences was clustered using CD-HIT [29], and cluster representatives were chosen at random from the resulting clusters. This resulted in a final set of around 8000 σ_2_ sequences and over 6500 σ_4_ sequences. This subsampled dataset was then used for domain-tree construction. Sequences were aligned using Muscle v5 and consequently trimmed using trimAl with a 0.02 cutoff parameter to reduce the phylogenetic noise in the data. A maximum-likelihood (ML)-based phylogenetic tree was constructed using IQ-TREE 3 [30], [31], with the best model chosen as Q.pfam+F+R6 [32] for domain 2 sequences and Q.pfam+R9 for domain 4 sequences using ModelFinder [33]. The rooting of the trees was done using the midpoint rooting method [34] and the minimal ancestor deviation (MAD) method [35]. All trees were visualised using iTOL [36].

## Results

To study the repertoire and evolution of σ-factors across bacteria and bacteriophages, we identified the various domains typically associated with σ-factors of the σ70 family using profile HMM searches of ∼1,400 genus-non-redundant bacterial genomes and over ∼350,000 bacteriophage genomes. Overall, we identified 28,909 bacterial and 32,488 bacteriophage proteins containing at least one of these domains.

Domains σ_2_ and σ_4_ recognise the bipartite bacterial promoter, with σ_4_ making the primary contact to the double-stranded ™35 element and σ_2_ binding the non-template strand of the unwound ™10 element [2]. Group 4, or extra-cytoplasmic function (ECF), proteins contain these two domains separated by a linker. Group 3 σ-factors contain an additional σ_3_. Finally, the major σ-factor, or Group 1 σ-factor, also contains σ_1_ (Figure 1A). σ_1_ contains two segments, and proteins lacking the N-terminal 1.1 domain are classified as group 2 σ-factors. For convenience, we refer to all proteins containing σ_1_ as group 1 σ-factors. We classified proteins with homology to these domains into the three σ-factor classes (∼35,000 proteins could be classified in this manner). Consistent with prior literature [37], [38], the most common σ-factor class in bacteria is group 4; this is true in bacteriophages as well. However, both bacteria and bacteriophages code for many proteins containing a single σ-factor domain -either σ_2_ or σ_4_. In fact, these single-domain proteins are the most common in bacteriophages and come second only to the group 4 σ-factors in bacteria. Thus, by taking a domain-level rather than a whole-protein-level approach to the detection of σ-factor-like proteins, we show that proteins, which would not typically be classified as σ-factors because they contain only a single domain - most commonly σ_4_ - are highly prevalent in both bacterial and bacteriophage genomes (Figure 1B and C).

**Figure 1.**
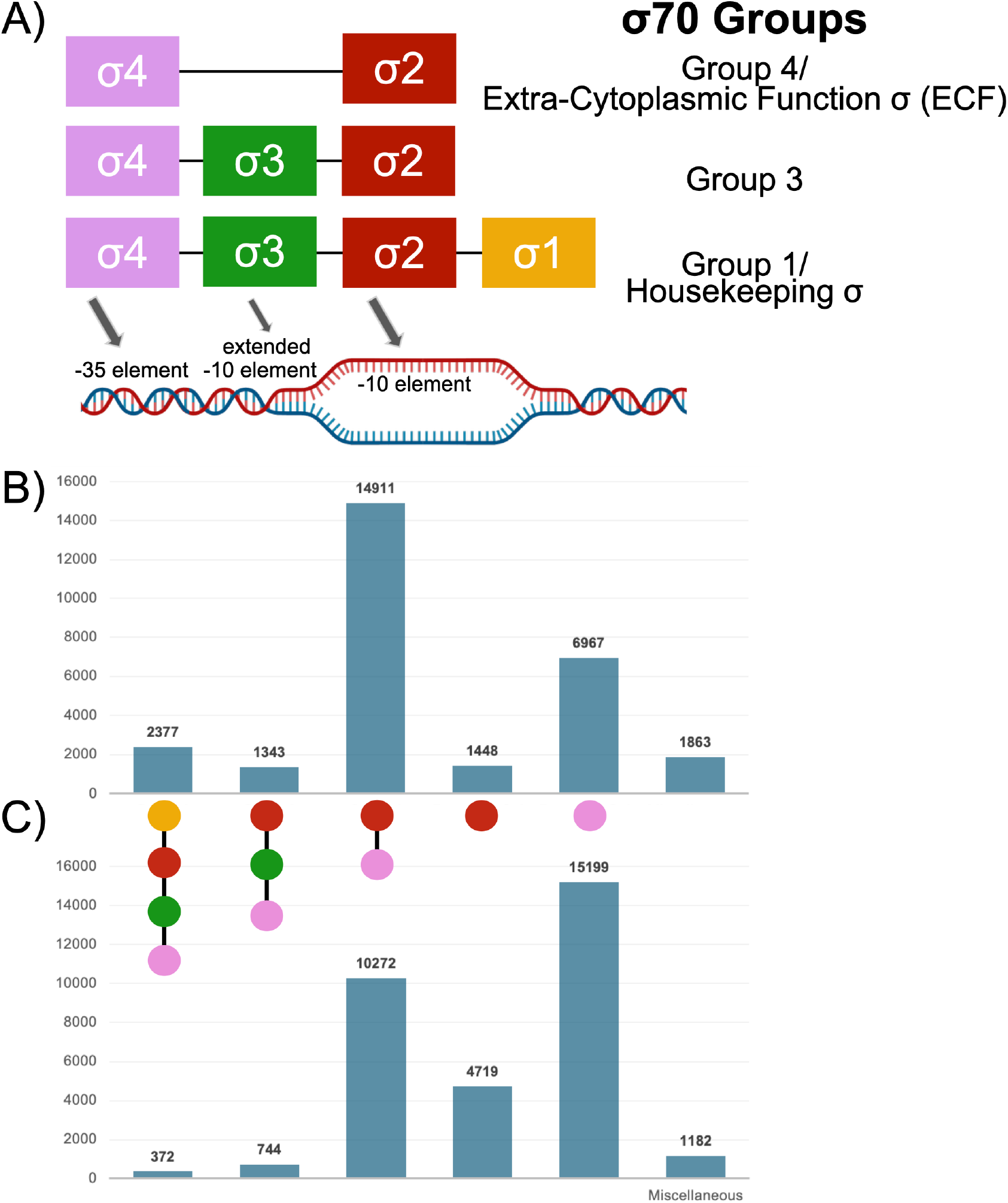
Distribution of Sigma Factor Proteins across bacteria and bacteriophages. (A) Schematic of σ70 domain architectures and the σ70 groups they define. Coloured boxes represent individual conserved domains (σ_1_, orange; σ_2_, dark red; σ_3_, green; σ_4_, lilac). Corresponding promoter elements recognised by each domain are shown on the DNA duplex. σ_2_ interacts with single-stranded DNA, and σ_4_ contacts the double-stranded DNA. The DNA duplex logo was made using BioRender. The number of proteins in bacteria, shown in (B), and bacteriophages, shown in (C); carrying domain combinations of sigma factor proteins, where each dot indicates the presence of a given domain (coloured as in panel A).

To investigate the evolutionary trajectory of σ-factors, we constructed maximum-likelihood, domain-level phylogenies for σ_2_ and σ_4_. Besides being the most prevalent in single-domain proteins, each domain also presents interfaces both to DNA and to the core RNAP. Further, the most basic domain architecture unit of σ-factors includes just these two domains. Under the idea that a protein domain is an independent folding and functional module, we assume that each domain can evolve independently via substitutions and short indels. Multiple domains, at any stage of their evolution, can fuse and dissociate to form different protein architectures with particular functional consequences. Though at a whole-protein level, different types of σ-factors could cluster separately as a result of their distinct domain architectures, domain-level phylogenies are independent of the effects of domain fusions and dissociations and help address the question of whether the evolution of individual domains has followed different trajectories in different σ-factor classes.

We aligned all σ_4_ sequences across bacteriophages and bacteria and across σ-factor classes and σ_4_-only proteins (a) freely using traditional multiple sequence alignments (using Muscle) and (b) constraining the alignment using the domain’s profile (using hmmalign). We built domain-level phylogenies for both alignments using FastTree against the complete set of over 45,000 (∼38,000 bacterial and ∼7,000 bacteriophage) σ_4_ sequences. Additionally, we also sampled over 6,500 sequences from the traditional MSA and constructed a phylogeny by choosing the best-fitting evolutionary model using IQ-TREE. We performed a similar analysis for the universal set of around 22,000 (∼19,000 bacterial and ∼3000 bacteriophage) σ_2_ sequences and a sampled set of around 8,000 sequences. We rooted these phylogenies using either the midpoint or the minimum ancestor deviation (MAD) approach. Here we describe our findings from the IQ-TREE-based phylogenies, rooted using the MAD approach. Similar results were obtained in the other trees as well, and these are shown in the supplementary material.

Rather than resolving into two monophyletic clades, bacteriophage-derived (inner ring, pink) and bacterial sequences (blue) are distributed across the full breadth of the tree. This patchy, non-monophyletic distribution holds at both the σ_2_ and σ_4_ levels (Figures 2A & 3A). This topology is consistent with multiple horizontal gene transfer events between bacteriophage and bacterial hosts involving genes encoding σ-factor domains. Within individual clades containing both bacterial and bacteriophage sequences, bacteriophage sequences disproportionately seem to branch earlier (Figures 2E & 3E). This is especially apparent in σ_4_ phylogenies but not to the same extent for σ_2_. This suggests that at least for σ_4_ sequences, a record of early branching sequences within various clades is retained among extant bacteriophage sequences rather than in bacterial ones.

**Figure 2.**
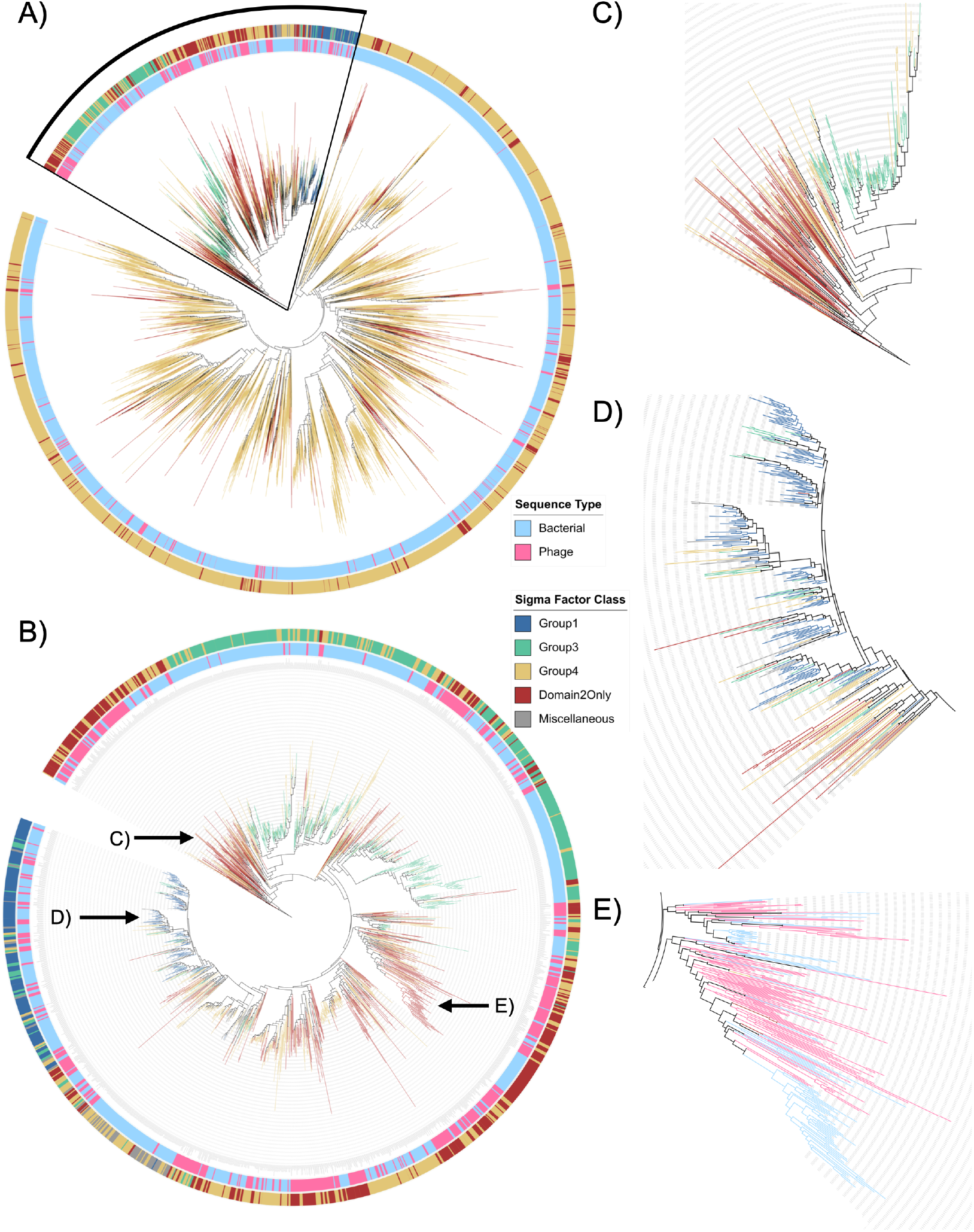
Maximum likelihood domain 2 phylogeny. (A) Minimal Ancestor Deviation (MAD) rooted domain 2 phylogenetic tree consisting of ∼8,000 σ_2_ sequences. The inner ring shows sequence type (bacterial, light blue; bacteriophage, pink); the outer ring is coloured by sigma factor class (Group1, blue; Group3, green; Group4, yellow; Domain2Only, red; Miscellaneous, grey). (B) Expanded view of the black pie in A. Arrows represent the zoomed-out regions. (C) Zoomed subclade showing early-emerging branches dominated by Group4 (yellow) and Domain2Only (red). (D) Zoomed subclade showing Group 1 (blue) forming a late-emerging, derived monophyletic cluster nested within the tree. (E) Zoomed region showing bacterial sequences (light blue) nested within predominantly phage sequences (pink).

**Figure 3.**
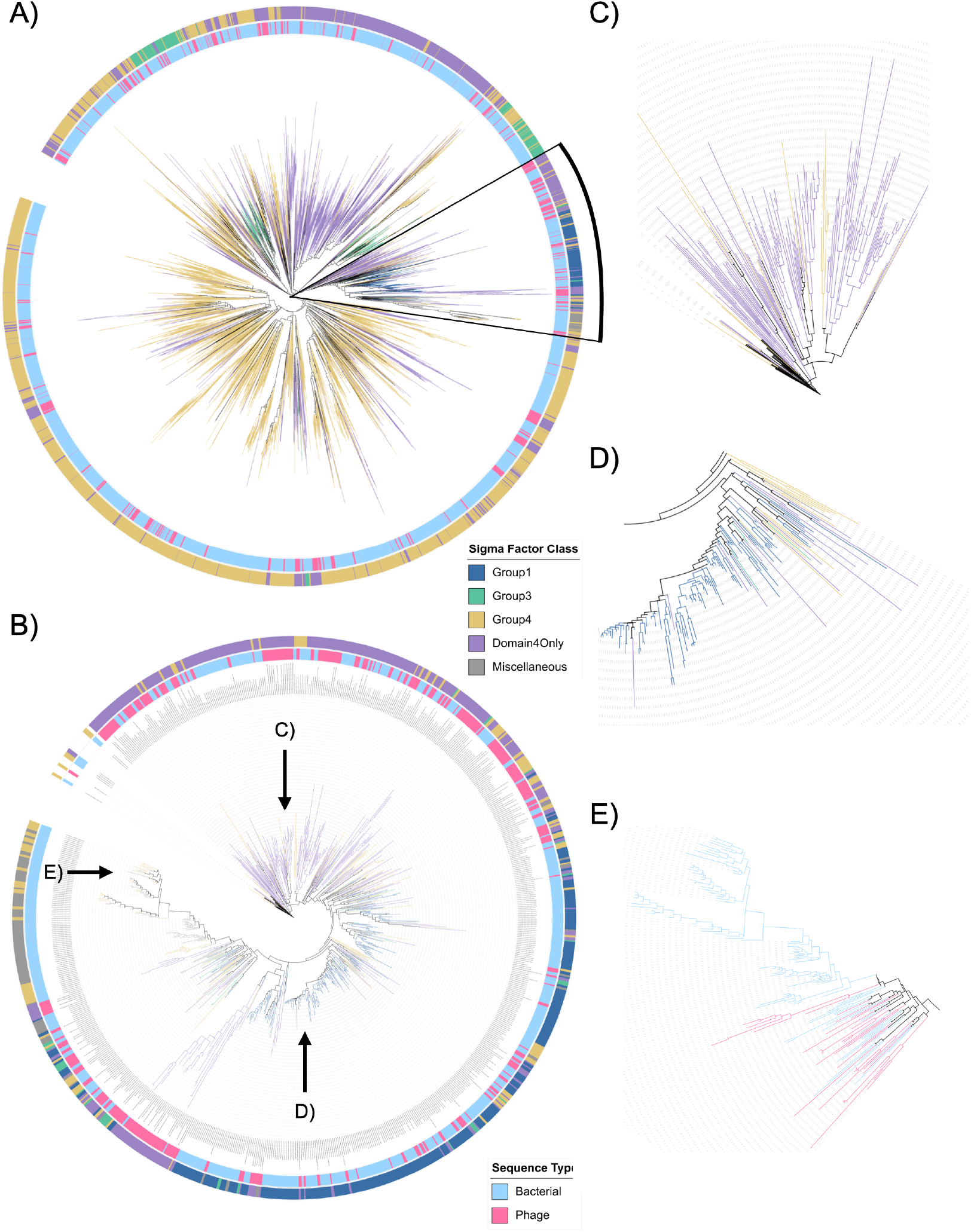
Maximum likelihood domain 4 phylogeny. (A) Minimal Ancestor Deviation (MAD) rooted domain 4 phylogenetic tree consisting of ∼6,500 σ_4_ sequences. The inner ring shows sequence type (bacterial, light blue; bacteriophage, pink); the outer ring is coloured by sigma factor class (Group1, blue; Group3, green; Group4, yellow; Domain4Only, lilac; Miscellaneous, grey). (B) Expanded view of the black pie in A. Arrows represent the zoomed-out regions. (C) Zoomed subclade showing early-emerging branches dominated by Group4 (yellow) and Domain4Only (lilac). (D) Zoomed subclade showing Group1 (blue) forming a late-emerging, derived monophyletic cluster nested within the tree. (E) Zoomed region showing bacterial sequences (light blue) nested within predominantly phage sequences (pink).

Group 1 and Group 3 σ-factors each form relatively coherent clusters; Group 1 forms a monophyletic clade in all trees but one. The only exception is in the HMM-aligned (PF04545) domain 4 phylogeny, where Group 1 has split into two clusters (Supplementary Figure 7B). Group 3 proteins consistently form two separate tight clusters. The sequence of these individual domains thus broadly seems to track their domain architecture-based classification. Group 4 sequences, however, depart from this pattern - rather than resolving as a single monophyletic arc, they are distributed across multiple, phylogenetically distant clades throughout the tree, which are often interspersed with the σ_4_-only or σ_2_-only proteins.

The patchy grouping of σ-factor sequences, especially those of the Group 4 and single-domain proteins, suggests that the evolution of these proteins has been shaped by a complex set of domain fusion and fission events. That said, however, certain clear patterns that are consistent across most of the phylogenies generated in this study emerge. Across all trees examined, Group 1 sequences, representing the primary, housekeeping class responsible for σ70-involved transcription, consistently form a late-diverging branch (Figures 2D & 3D). This implies that the housekeeping σ70 architecture, despite its near-universal conservation and central role in bacterial transcription today, might not be the ancestral σ-factor form. Group 3 σ-factors seem to also be late-diverging but do not share any immediate common ancestor with the Group 1 proteins (Figure 2C). Along paths leading ultimately to the emergence and divergence of the monophyletic group 1 σ-factors or to group 3 proteins, single-domain proteins occupy the earliest-diverging positions alongside some group 4 sequences (Figures 2C & 3C), a pattern that is consistent across traditional and constrained alignment methods as well as MAD and midpoint rooting (see supplementary Fig. 2-8). We also note here that these earliest branching lineages include a large number of bacteriophage-derived sequences.

Taken together, our observations suggest a model in which minimal, single-domain σ-factor modules, comprising domain 2 and/or domain 4 alone, without the additional regulatory regions present in full-length sigma factors, represent an ancestral state from which more structurally elaborate, multi-domain sigma factors, including the housekeeping major σ-factor, subsequently evolved. That said, however, the evolution of σ-factor sequences is highly complex and probably has also involved minimisation in which many single-domain proteins have derived from multi-domain σ-factors.

## Discussion

Our study suggests that σ-factors evolved in a stepwise manner by domain fusion or accretion, starting from single-domain proteins capable of binding both DNA and RNAP (Supplementary Figures 9 & 10), which came together to form the minimal σ-factor architecture characterising the ECF σ-factor; other domains, which seem to exist rarely in isolation in extant bacterial or bacteriophage genomes (Supplementary Figure 1), then fused to the minimal σ-factor architecture to form more elaborate forms with complex control of DNA binding specificity (Figure 4). The sequence record of these ancestral single-domain proteins appears to be better preserved in bacteriophage genomes rather than in bacterial ones.

**Figure 4.**
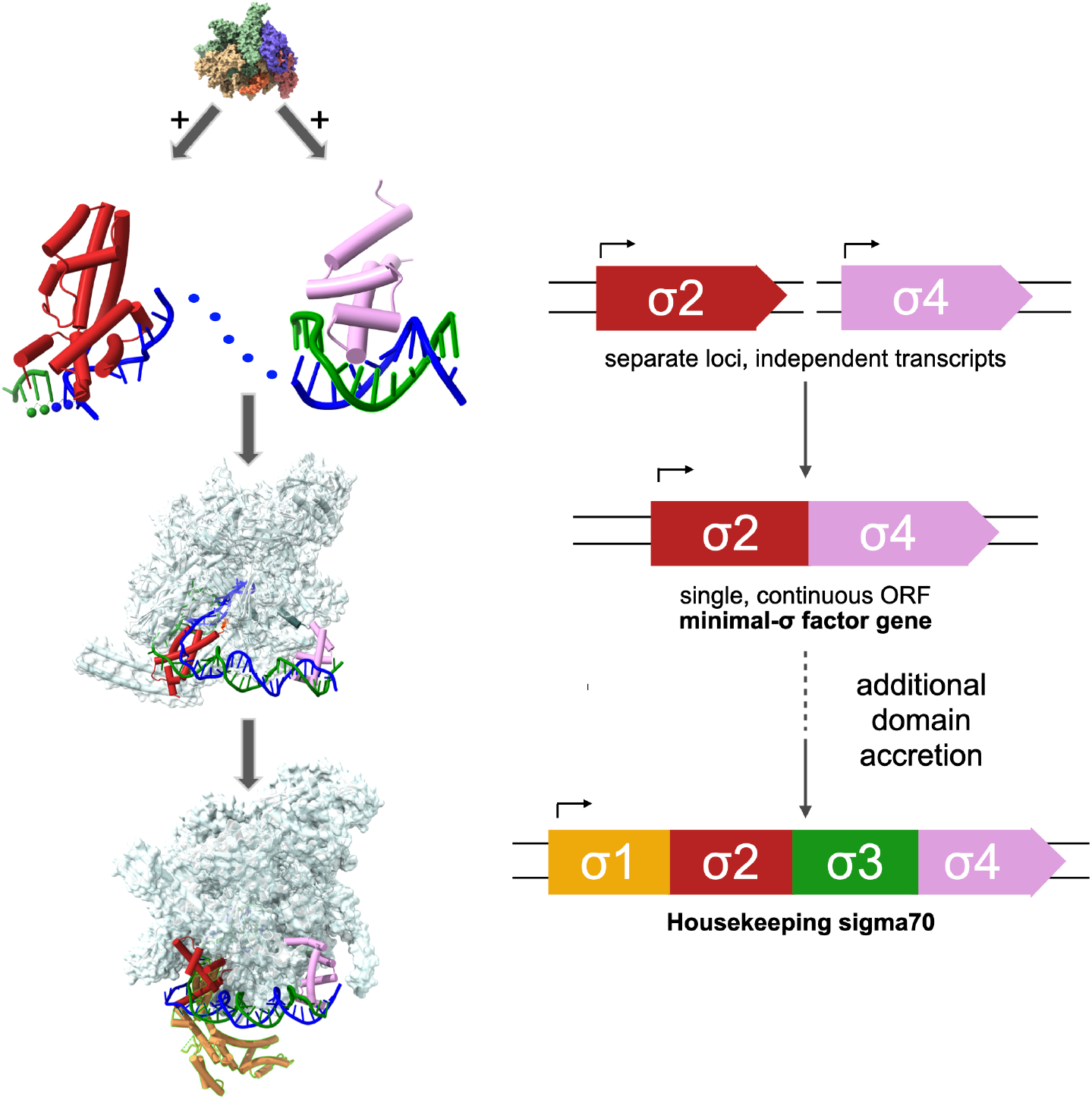
Model for the evolution of sigma factor proteins. (Left) Structural model illustrating how individual σ domains engage promoter DNA. The σ_4_ domain (lilac) binds double-stranded promoter DNA (blue/green duplex), while the σ_2_ domain (red) binds single-stranded, melted DNA (blue/green single strands). The RNAP core enzyme (top, multicoloured surface) associates with these two σ domains. Downstream, the minimal ECF-type σ-factor (lower-middle structure) and the full housekeeping σ70 factor (bottom structure) are shown docked onto the RNAP core with DNA, illustrating assembly of the transcription-initiation complex. (Right) Proposed evolutionary model for domain/gene fusion giving rise to modern σ-factors. σ_2_ and σ_4_ are initially encoded as separate genes at independent loci, each with its own transcript (top). Fusion of these genes into a single, continuous open reading frame produces a minimal σ-factor gene (σ_2_–σ_4_), corresponding to the ECF-type σ-factor. Subsequent accretion of additional domains (σ_3_, σ_1_) onto this minimal gene gives rise to the more elaborate housekeeping σ70 gene architecture.

Scenarios on the evolution of the RNAP suggest that elongation evolved first [39]. The minimal form of RNAP (comprising the catalytic and the assembly subunits of the bacterial RNAP) is conserved in bacteria as well as in the archaea and eukaryotes, which share additional common components [40]. Proteins that help in transcription elongation are conserved across domains of life. In contrast, initiation factors differ between the bacterial and the archaeal-eukaryotic RNAP. σ-factors in bacteria form a holoenzyme complex with the core RNAP, which then finds and binds to the promoter. However, the prior formation of a complex of RNAP and its specificity-determining components is not a prerequisite for transcription initiation by extant multi-subunit RNAP. Initiation complex components of archaeal and eukaryotic RNAP can first assemble on the promoter or enhancer DNA and then recruit the enzyme [39]. Even among σ-factors, the mechanism by which the binding of these proteins to DNA in the absence of the RNAP is inhibited has been fleshed out for the major, housekeeping sigma factors; the domains responsible for this inhibition are absent in the more minimal ECFs [41]. Further, there is some evidence that the major σ-factor in *Bacillus subtilis* can directly bind part of the promoter DNA [42]. The evolution of the relationship between the RNAP and its initiation factors in eukaryotes is especially complex - some factors that are dissociable in the mRNA-transcribing RNAP-II are fused to the main components of RNAP in RNAP-I and RNAP-III, which transcribe translation-associated RNA genes [40], [43]. Finally, it has been argued that the DNA-binding HTH motif of σ-factors may be homologous to that in the archaeo-eukaryal TFIIB; the evidence is much stronger for σ_4_ than for σ_2_ [44] (Supplementary Figure 12). This suggests a deep common origin for specificity-determining factors for both bacterial and archaeo-eukaryal multi-subunit RNAPs.

Though it is intuitive to posit that basal regulators of transcription, such as σ-factors and archaeal-eukaryotic transcription initiation factors, being part of the initiating RNAP complex, are evolutionarily older than specific TFs that activate or repress the expression of small subsets of genes, this is by no means a given. Sequence analysis performed two decades ago suggested that some protein sequence families linked to specific TFs are more prevalent across the tree of life than σ-factors and could therefore be older [45]. More recent phylogenetic reconstruction studies based on orthologous groups of TFs also indicated that the common ancestor of the winged-helix domain superfamily, commonly found in specific TFs, might trace back to a time point prior to the last universal common ancestor, whereas σ-factors may go back to the last bacterial common ancestor [46]. Further, first principles of protein evolution suggest that complex sequences evolved from simpler motifs by duplication and further evolution [47], reviewed in a perspective by Romero, Rabin and Tawfik in 2016 [48]. Fusion of simple domains to form more complex protein architectures also has a role to play [49], [50]. If the HTH motif of σ-factors and archaeal and eukaryotic initiation factors share a common ancestor, then it suggests diversification of this ancestral sequence into two lineages, one of which then underwent a series of fusion events to eventually evolve the housekeeping major σ-factor. The evolution of the major σ-factor might have also involved duplications of shorter fragments [51].

A caveat of our study arises from the robustness of alignments, trees, and rootings. We have addressed these using multiple approaches at each level and identified aspects that are consistent across (Supplementary Figures 2-8). We have also subsampled sequences while maintaining overall sequence diversity and shown that this does not materially change the take-home messages. σ_4_ sequences are much shorter than domain 2 sequences, and that can create greater uncertainty in the phylogenies of the former; this may be reflected in the greater variation in tree topologies that we seem to observe across methods for domain 4 alignments. Of particular concern is always where the root is positioned. Mid-point rooting assumes constant evolutionary rates, an assumption that is seldom met. Nevertheless, there is no reason to suspect that early branching lineages in most of our trees show unusually long root-tip distances. However, we have also used the MAD method, which is designed to be robust to the presence of heterotachy, to root our trees and find that the major findings hold [35]. We also aligned an archaeal TFIIB (PDB ID: 1D3U) to our existing σ_4_ alignment (∼6,500 sampled sequences) and built a FastTree-based phylogeny using TFIIB as the outgroup. Though the overall topology of the tree is different from the other trees we have generated, group 1 and group 3 sequences still emerge as late-diverging in this tree as well (Supplementary Figure 11). Note that since TFIIB is a very distant relative, this alignment should be considered as adding weaker supporting evidence.

In an ancestral cell, RNAP possibly bound DNA non-specifically and initiated transcription randomly. Some DNA-binding proteins, acting as repressors, might have simply blocked its binding at some sites by steric hindrance. Other proteins, having evolved an interface to recruit the RNAP, acted as activators, making some sites on the DNA more attractive for the RNAP to bind to and initiate transcription. Some of these proteins were probably σ_4_-only or σ_2_-only proteins. There are structures of individual σ_4_ and σ_2_ domains extracted from a sigma factor protein bound to double-stranded (PDB ID: 2H27, [52]) and single-stranded DNA (PDB ID: 3UGO, [53]), suggesting that these domains can bind to DNA in the absence of the RNAP. The binding sites of the two proteins on the DNA might have been close together in a form similar to the bipartite promoter architecture that is familiar to us today. This is not all that improbable: bacterial promoter sequences can evolve from both random DNA sequences [54] as well as be found in numbers within A+T-rich protein-coding gene sequences [55]. A fortuitous fusion of genes encoding these two single-domain proteins would have been selected for by persistent proximity of the two promoter elements on the ancient DNA. Assuming that σ_2_ does not on its own make any initial contact with double-stranded DNA in or around promoter-like sequences, the difficult recognition and binding of transiently single-stranded/base-flipped DNA by σ_2_ [53], [56], would have been enabled by the reduced degree of freedom offered by its fusion to σ_4_. Finally, because both domains contact RNAP, the fusion protein - the common ancestor of the most minimal σ-factors - would have possessed an increased ability to bind the RNAP as well by presenting a larger protein-protein interaction interface. Preferential binding of this protein to the RNAP through this large interface before it could bind DNA would have given the RNAP its holoenzyme form and specificity in promoter recognition. The linker between the two domains in ECF proteins is weakly conserved in sequence, but affords additional interactions with the RNA exit channel [57], a function which could be performed by a newly acquired domain 3.2 in more elaborate σ-factors [58]. Finally, the selectivity filtering function offered by another domain (domain 1.2) provided RNAP the ability to tolerate more variations in promoter sequence and architecture and emerge as a global basal TF that is responsible for transcription of housekeeping genes across bacteria [59].

## Acknowledgements

We thank Madhumitha Krishnaswamy and Sarthak Ghatkar for curating the bacterial and bacteriophage genomes used in this study. We thank Ashitha Arun, B. Gopal, Ganesh Muthu, Inder Raj Singh and Sunil Laxman for comments on the paper. This work was funded by Department of Atomic Energy, Government of India, project identification no. RTI 4006 and RTI 4018.

## Data Availability

Data are available in Figshare (https://doi.org/10.6084/m9.figshare.34219659). Supplementary Figures 1–12 are provided as Supplementary Material.

## AI Use Declaration

Claude Opus 5.5 (Anthropic) was used to find specific literature and to clean up and format text. All suggestions made by AI were checked by the authors.

