## Supplementary Figures 1-12 for "Phylogenetic evidence for the emergence of bacterial σ-factors by domain fusion"

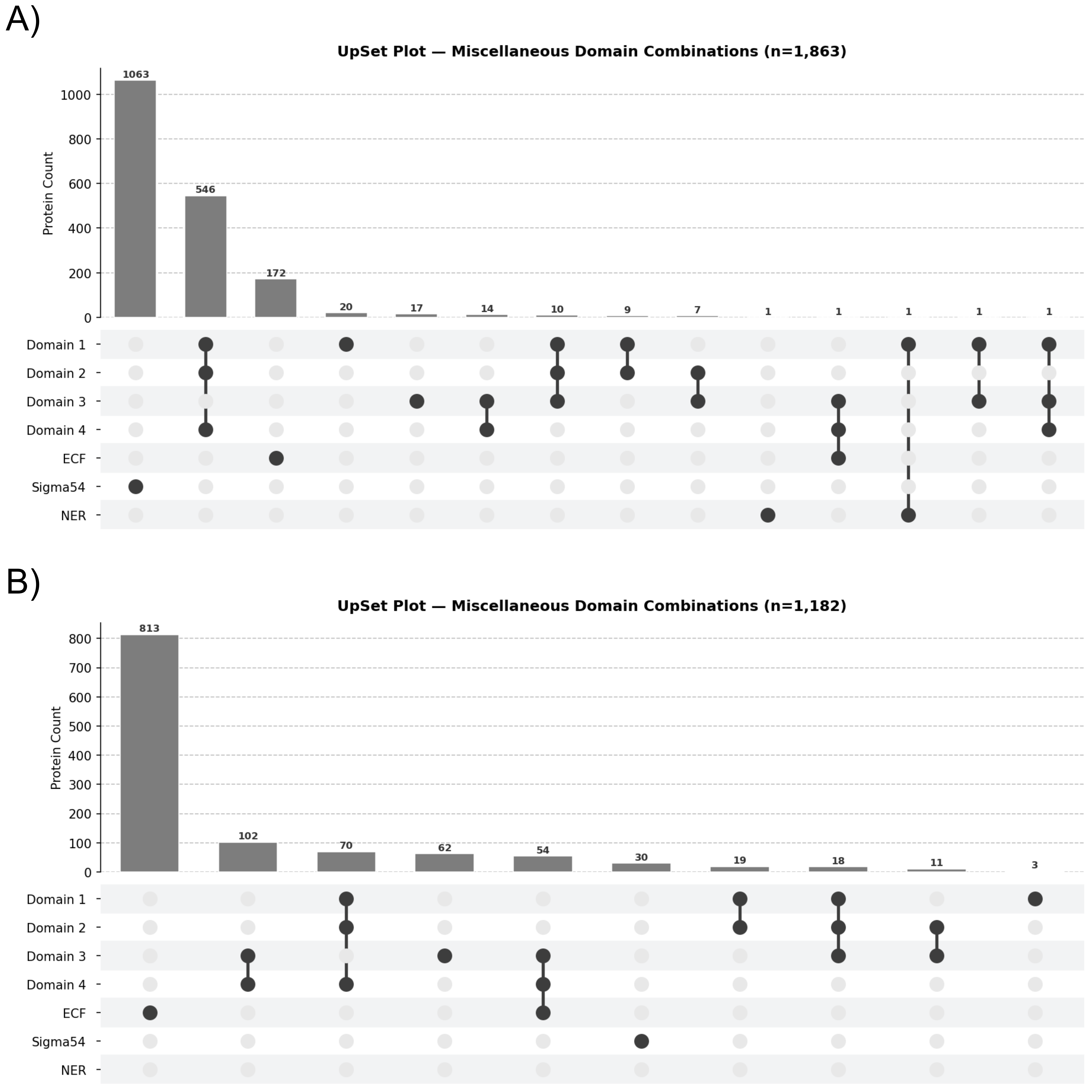


**Supplementary Fig. 1.** Domain composition of miscellaneous sigma factor hits. UpSet plots showing combinations of conserved domains (Domain 1-4, ECF, Sigma54, NER) identified in proteins that did not classify into the main sigma factor groups, for (A) bacterial (n=1,863) and (B) phage (n=1,182) proteins. Bar height indicates the number of proteins containing a given domain combination (dot matrix below), sorted by frequency.

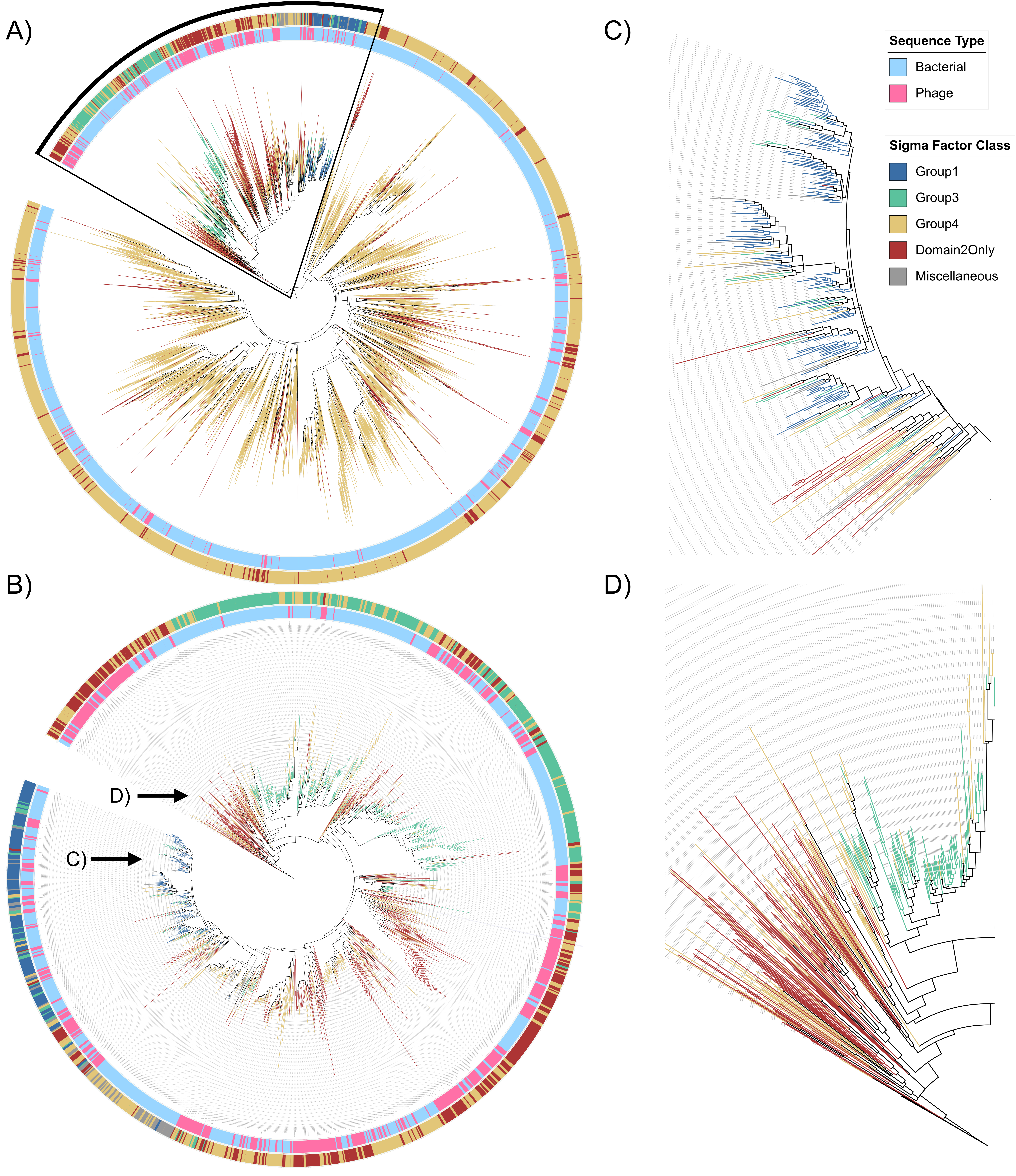


**Supplementary Fig. 2.** Maximum likelihood domain 2 phylogeny visualized using iTOL. (A) Midpoint-rooted, MUSCLE-aligned domain 2 phylogenetic tree consisting of over 8,000 σ2 sequences. The inner ring shows sequence type (bacterial, light blue; bacteriophage, pink); the outer ring is colored by sigma factor class (Group1, blue; Group3, green; Group4, yellow; Domain2Only, red; Miscellaneous, grey). (B) Expanded view of the black pie in A. Arrows represent the zoomed-out regions. (C) Zoomed subclade showing Group 1 (blue) forming a late-emerging, derived monophyletic cluster nested within the tree. (D) Zoomed subclade showing early-emerging branches dominated by Group4 (yellow) and Domain2Only (red).

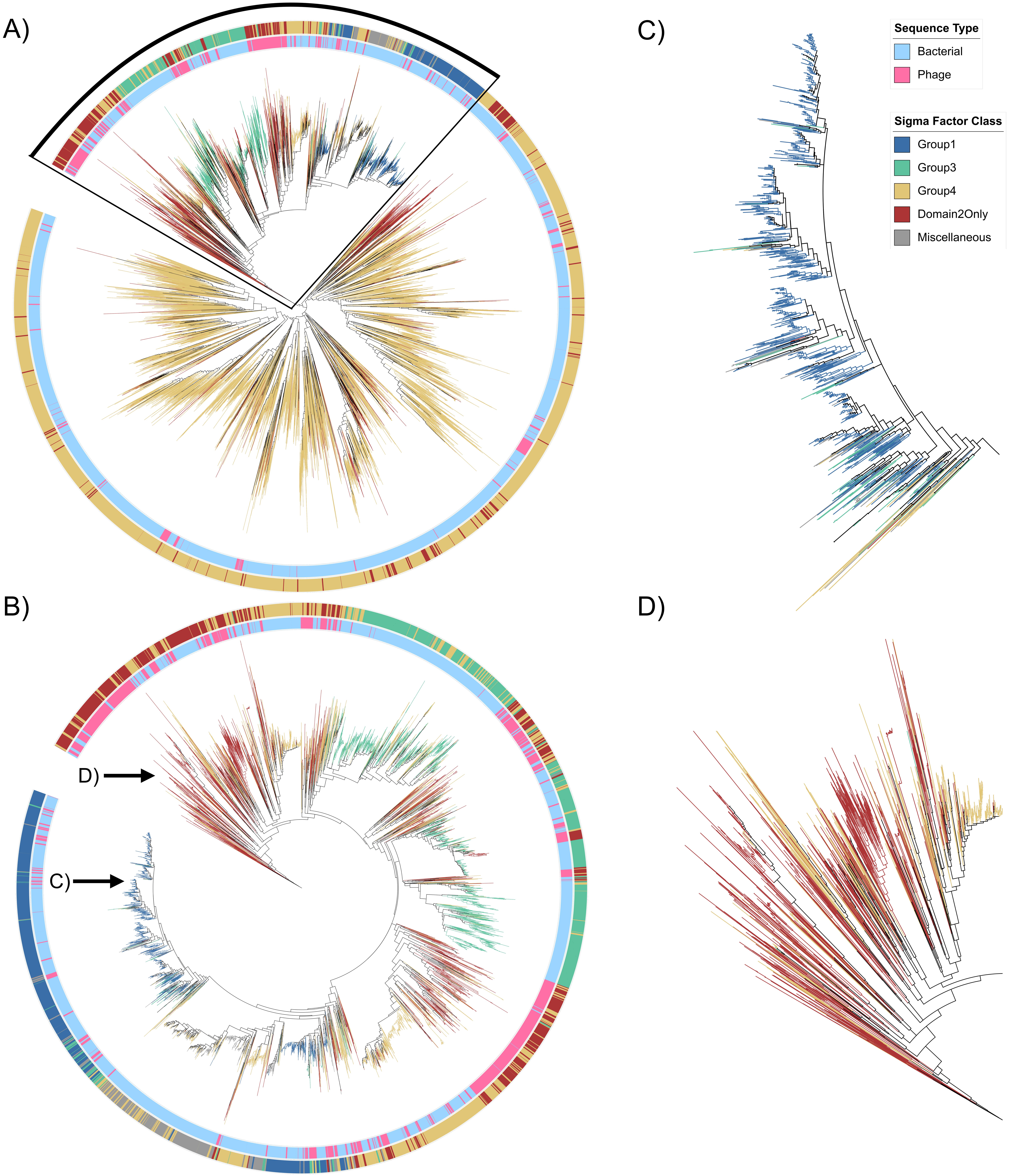


**Supplementary Fig. 3.** Maximum likelihood domain 2 phylogeny visualized using iTOL. (A) Midpoint-rooted, HMM-aligned domain 2 phylogenetic tree consisting of over 22,000 σ2 sequences. The inner ring shows sequence type (bacterial, light blue; bacteriophage, pink); the outer ring is colored by sigma factor class (Group1, blue; Group3, green; Group4, yellow; Domain2Only, red; Miscellaneous, grey). (B) Expanded view of the black pie in A. Arrows represent the zoomed-out regions. (C) Zoomed subclade showing Group 1 (blue) forming a late-emerging, derived monophyletic cluster nested within the tree. (D) Zoomed subclade showing early-emerging branches dominated by Group4 (yellow) and Domain2Only (red).

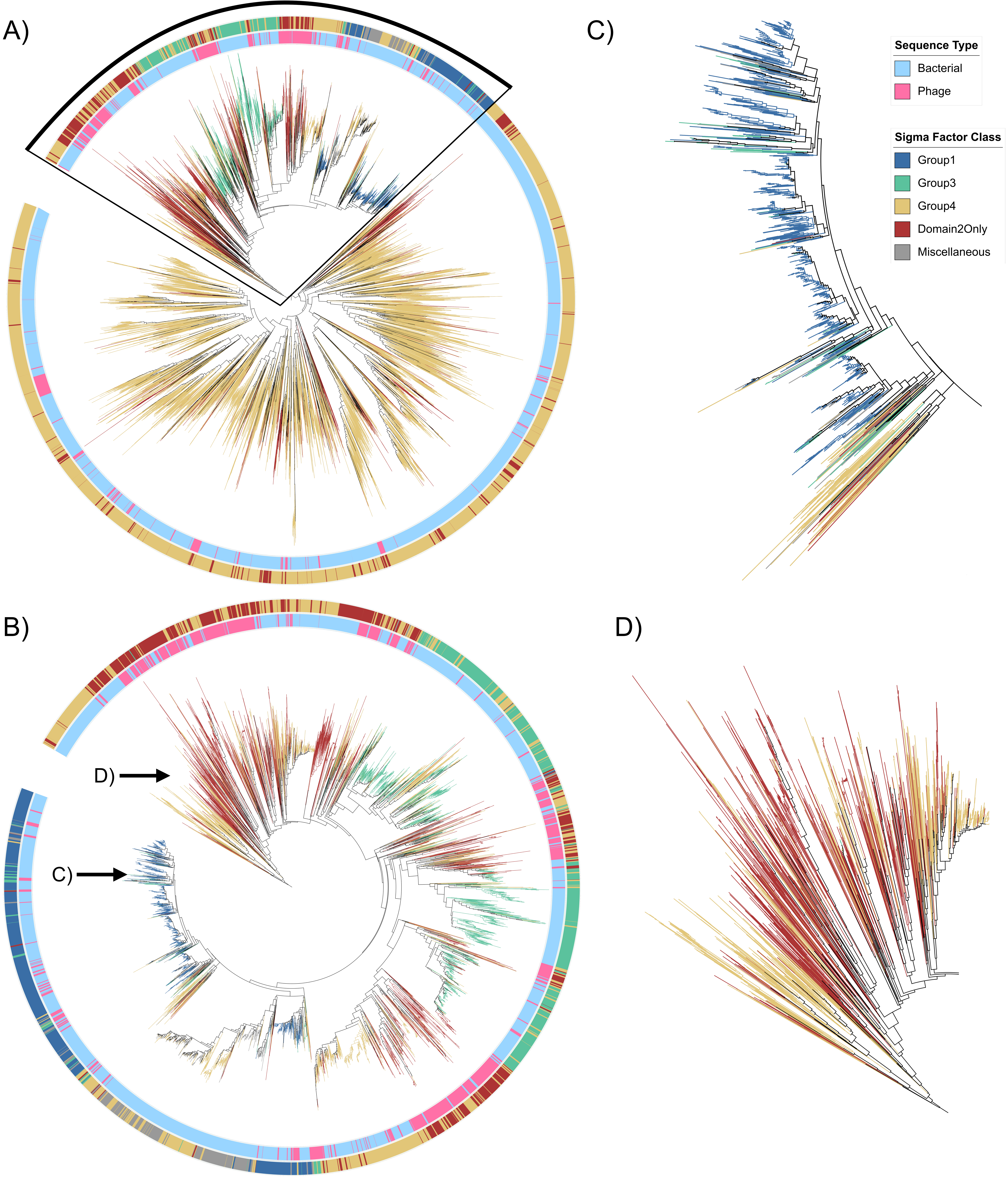


**Supplementary Fig. 4.** Maximum likelihood domain 2 phylogeny visualized using iTOL. (A) Midpoint-rooted, MUSCLE-aligned domain 2 phylogenetic tree consisting of over 22,000 σ2 sequences. The inner ring shows sequence type (bacterial, light blue; bacteriophage, pink); the outer ring is colored by sigma factor class (Group1, blue; Group3, green; Group4, yellow; Domain2Only, red; Miscellaneous, grey). (B) Expanded view of the black pie in A. Arrows represent the zoomed-out regions. (C) Zoomed subclade showing Group 1 (blue) forming a late-emerging, derived monophyletic cluster nested within the tree. (D) Zoomed subclade showing early-emerging branches dominated by Group4 (yellow) and Domain2Only (red).

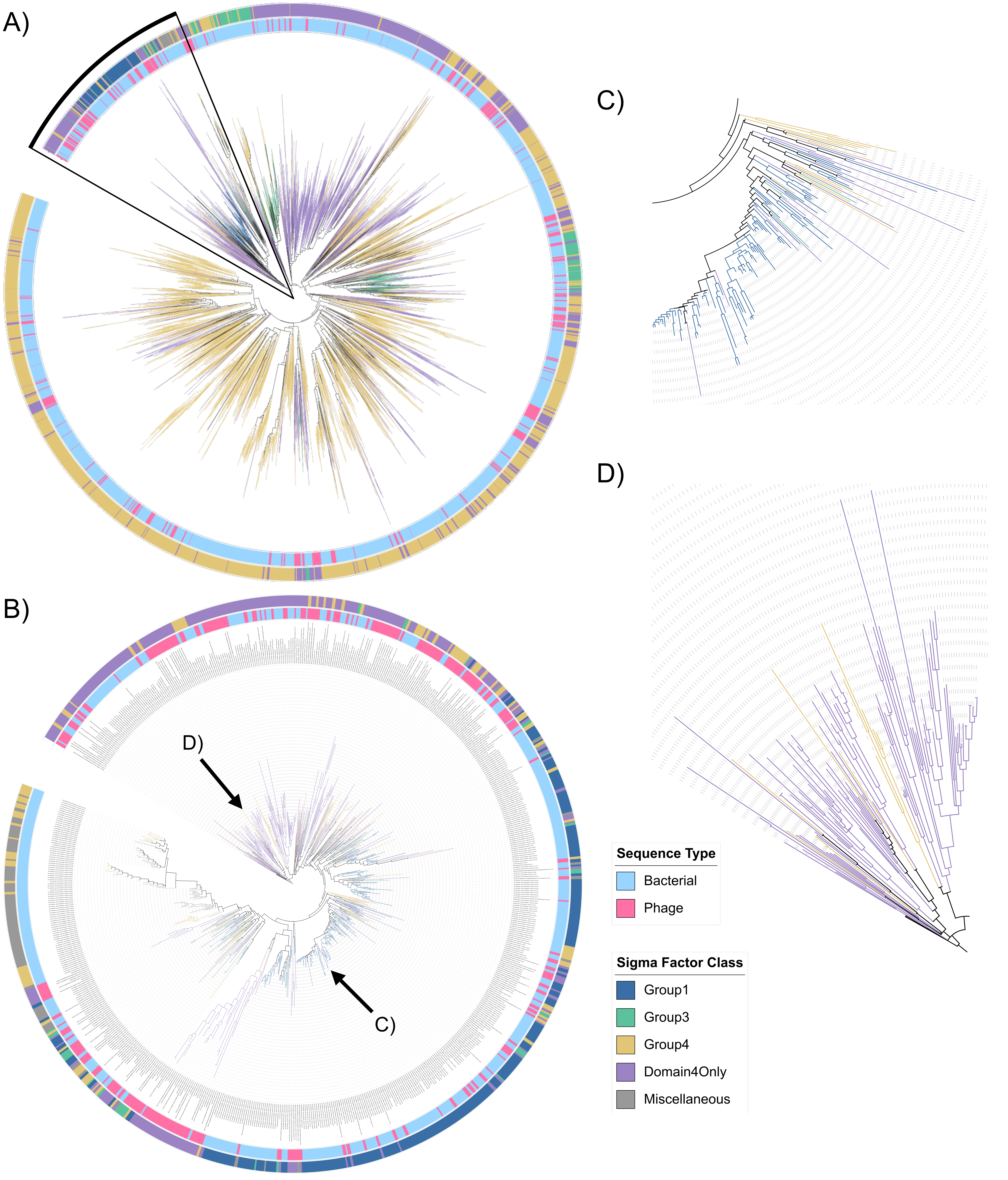


**Supplementary Fig. 5.** Maximum likelihood domain 4 phylogeny visualized using iTOL. (A) Midpoint-rooted, MUSCLE-aligned domain 4 phylogenetic tree consisting of over 6500 σ4 sequences. The inner ring shows sequence type (bacterial, light blue; bacteriophage, pink); the outer ring is coloured by sigma factor class (Group1, blue; Group3, green; Group4, yellow; Domain4Only, lilac; Miscellaneous, grey). (B) Expanded view of the black pie in A. Arrows represent the zoomed-out regions. (C) Zoomed subclade showing Group1 (blue) forming a late-emerging, derived monophyletic cluster nested within the tree. (D) Zoomed subclade showing early-emerging branches dominated by Group4 (yellow) and Domain4Only (lilac).

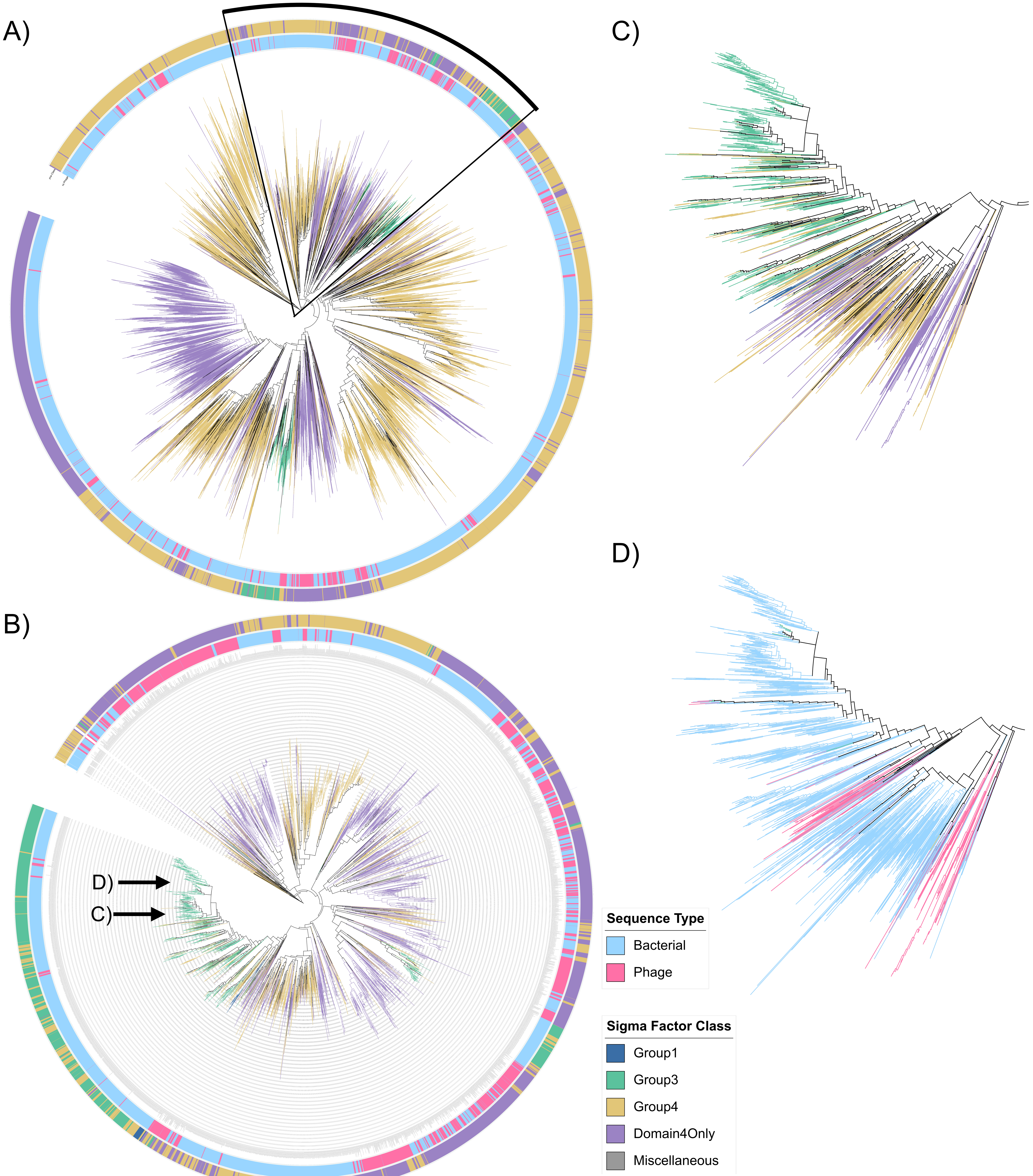


**Supplementary Fig. 6.** Maximum likelihood domain 4 phylogeny visualized using iTOL. (A) Midpoint-rooted, HMM-aligned (with profile PF08281) domain 4 phylogenetic tree consisting of over 24,000 σ4 sequences. The profile PF08281 seems to have a bias towards picking the Domain4Only, Group3, Group4 proteins only; it picked very few Group1 proteins. The inner ring shows sequence type (bacterial, light blue; bacteriophage, pink); the outer ring is coloured by sigma factor class (Group1, blue; Group3, green; Group4, yellow; Domain4Only, lilac; Miscellaneous, grey). (B) Expanded view of the black pie in A. Arrows represent the zoomed-out regions. (C) Zoomed subclade showing early-emerging branches dominated by Group4 (yellow), Domain4Only (lilac); and Group 3 being a late-emerging, derived clade. (D) Zoomed region showing bacterial sequences (light blue) nested within predominantly phage sequences (pink).

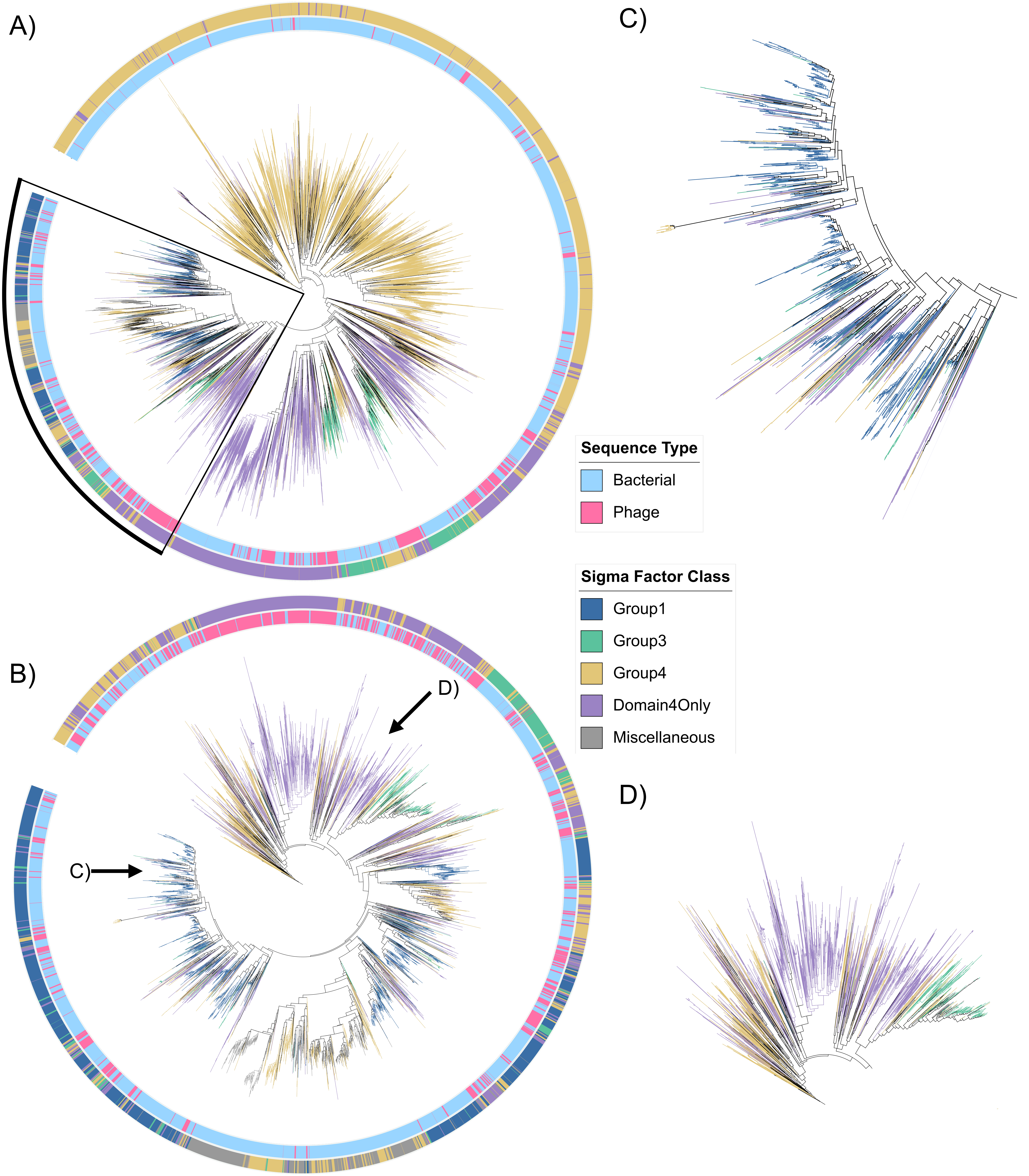


**Supplementary Fig. 7.** Maximum likelihood domain 4 phylogeny visualized using iTOL. (A) Midpoint-rooted, HMM-aligned (with profile PF04545) domain 4 phylogenetic tree consisting of over 21,000 σ4 sequences. The profile PF04545 picked all sigma factor proteins. The inner ring shows sequence type (bacterial, light blue; bacteriophage, pink); the outer ring is coloured by sigma factor class (Group1, blue; Group3, green; Group4, yellow; Domain4Only, lilac; Miscellaneous, grey). (B) Expanded view of the black pie in A. Arrows represent the zoomed-out regions. (C) Zoomed subclade showing Group1 as two separate but late-emerging clusters. (D) Zoomed subclade showing early-emerging branches dominated by Group4 (yellow), Domain4Only (lilac); and Group 3 being a late-emerging, derived clade.

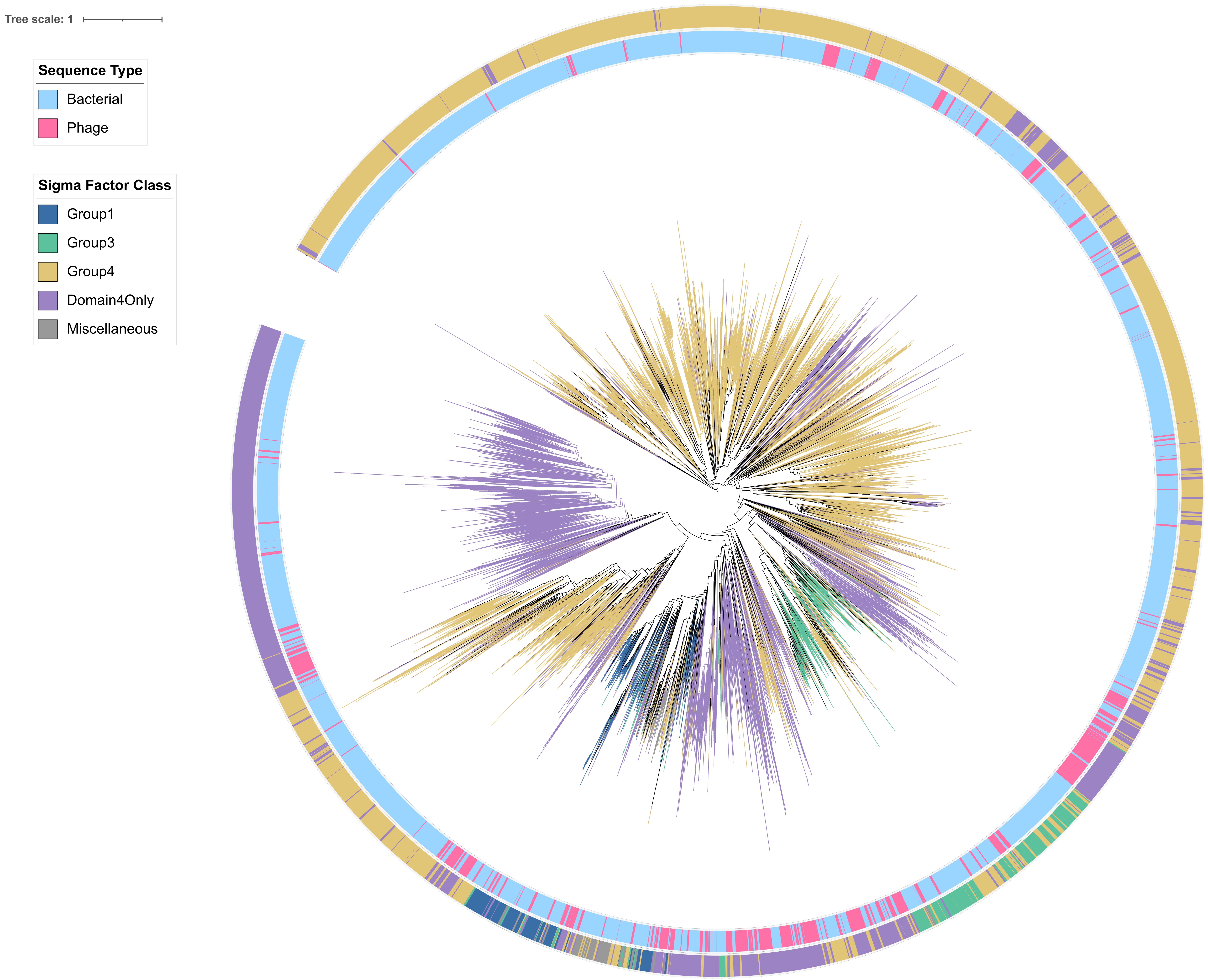


**Supplementary Fig. 8.** Maximum likelihood domain 4 phylogeny, visualized using iTOL. (A) Midpoint-rooted, MUSCLE-aligned domain 4 phylogenetic tree consisting of over 45,000 σ4 sequences. The inner ring shows sequence type (bacterial, light blue; bacteriophage, pink); the outer ring is coloured by sigma factor class (Group1, blue; Group3, green; Group4, yellow; Domain4Only, lilac; Miscellaneous, grey).

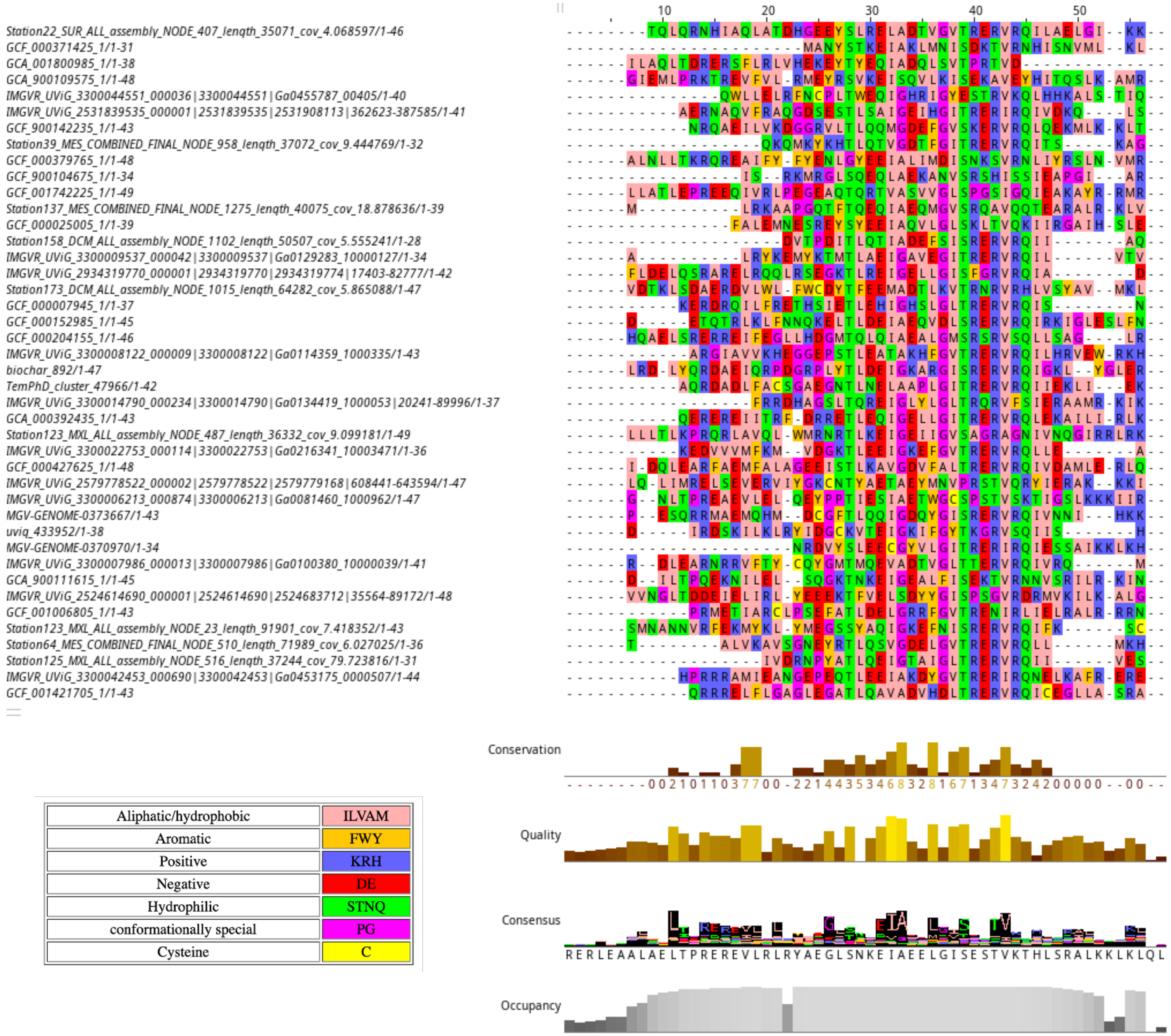


| Aliphatic/hydrophobic | ILVAM |
|---|---|
| Aromatic | FWY |
| Positive | KRH |
| Negative | DE |
| Hydrophilic | STNQ |
| conformationally special | PG |
| Cysteine | C |

**Supplementary Fig. 9.** Multiple sequence alignment of domain 4-only proteins. Amino acid residues are colored based on their physicochemical properties. Positively charged amino acids conserved (K and R) towards the C-terminal region of domain 4 involved in promoter binding.

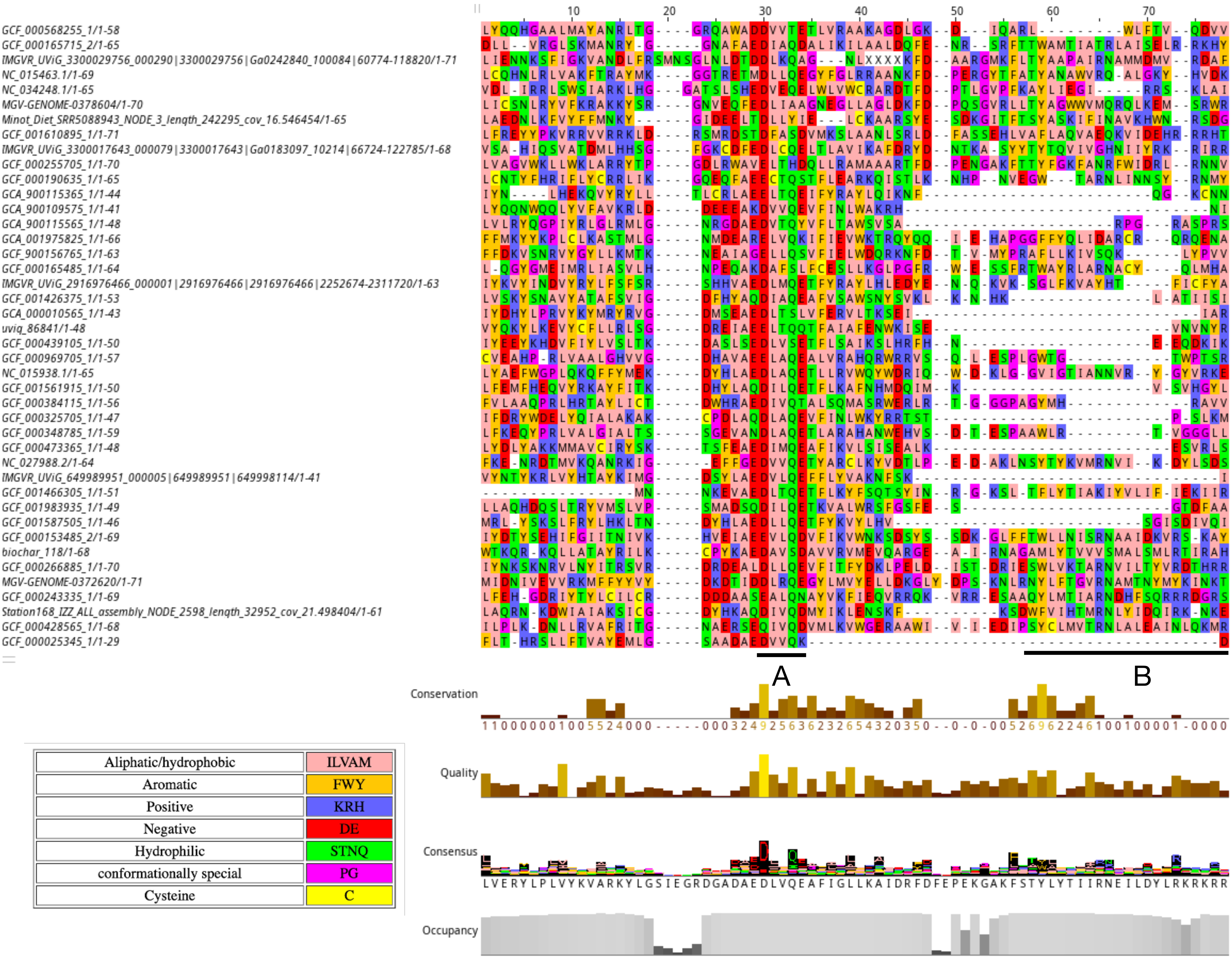


**Supplementary Fig. 10.** Multiple sequence alignment of domain 2-only proteins. Amino acid residues are colored based on their physicochemical properties. The domain 2 region 2.2 consisting of residues D, Q and E (patch A) involved in RNAP binding, while the region 2.4 consisting of aromatic amino acids (F, W, Y) and R (patch B) involved in promoter interaction.

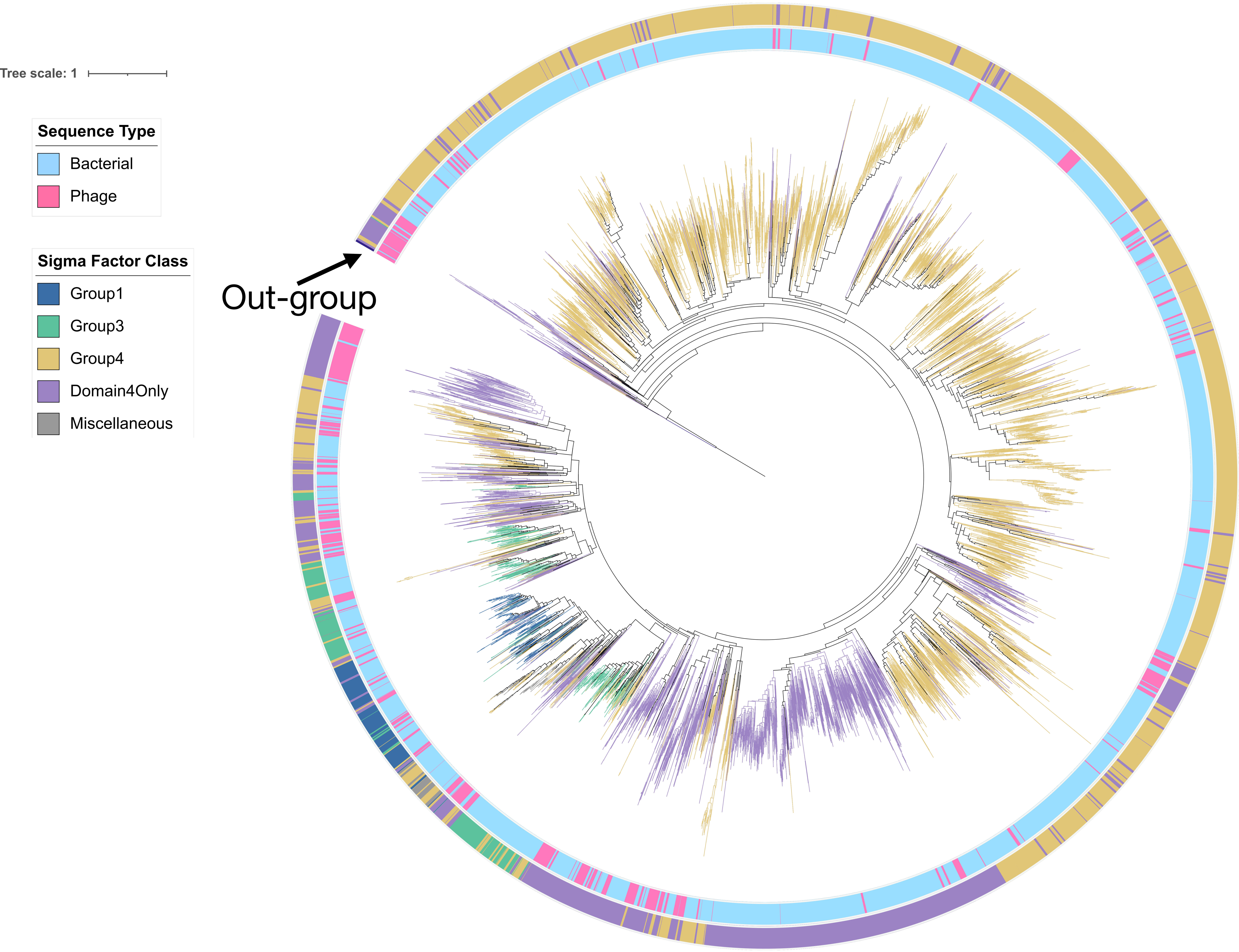


**Supplementary Fig. 11.** Domain 4 tree with Transcription Factor (II)B of *Pyrococcus woesei* as the outgroup (visualized using iTOL). The inner ring shows sequence type (bacterial, light blue; bacteriophage, pink); the outer ring is coloured by sigma factor class (Group1, blue; Group3, green; Group4, yellow; Domain4Only, lilac; Miscellaneous, grey; Outgroup, black).

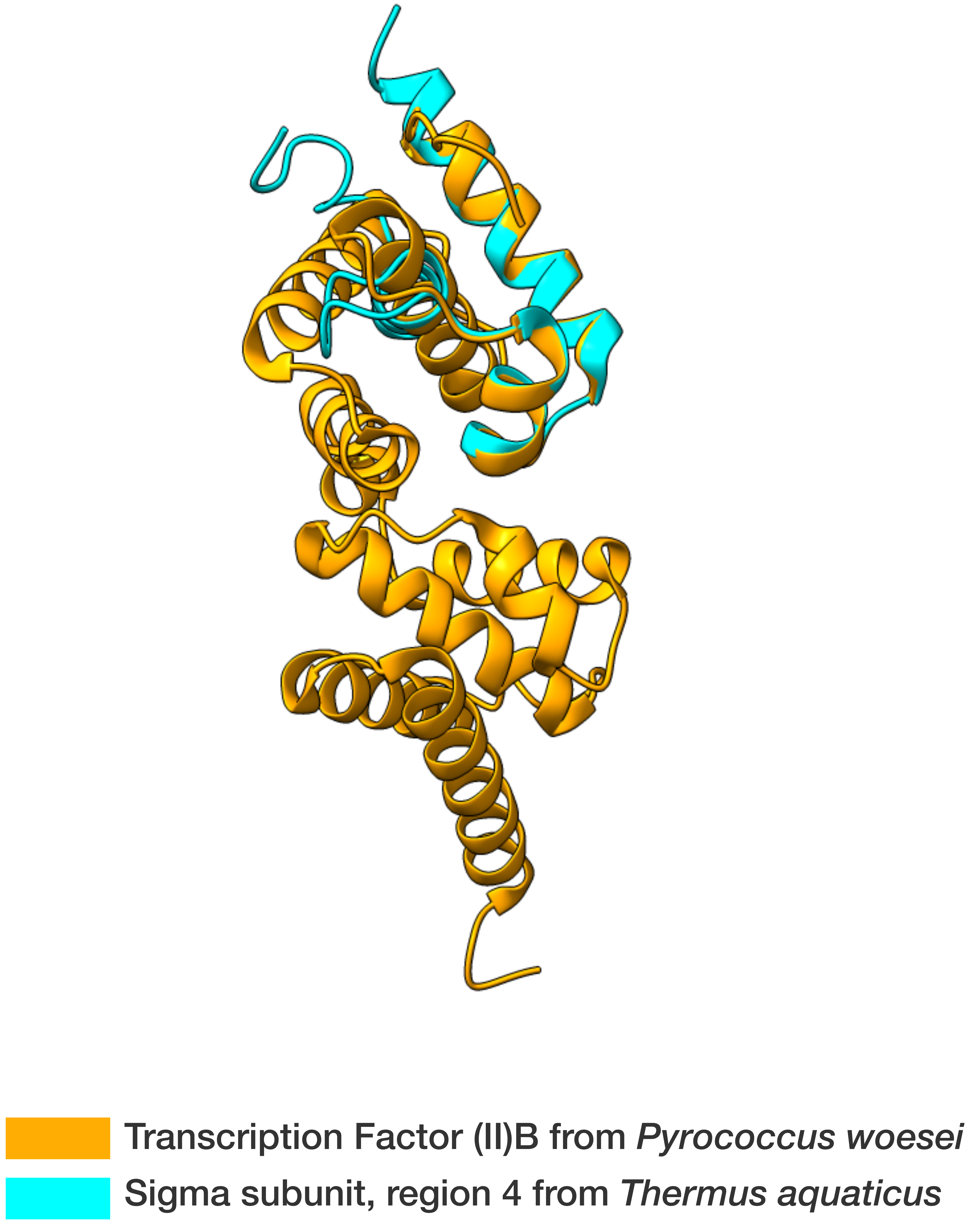


**Supplementary Fig. 12.** Structural alignment of Transcription Factor (II)B from *Pyrococcus woesei* and sigma-subunit region 4 from *Thermus aquaticus.* The RMSD of 31 pruned atom pairs of the helix-turn-helix of region 4 of the sigma subunit is 0.751 angstroms (across all 61 pairs, RMSD is 6.940).
